# Macrophages as Key Mediators of Trp-1-Specific CD4^+^ T Cell Therapy for Murine Melanoma

**DOI:** 10.1101/2023.12.20.572715

**Authors:** Adam Q. He, Dawei Zou, Wenhao Chen

## Abstract

There is a growing body of evidence indicating that CD4^+^ T cells, alongside CD8^+^ T cells, can effectively combat cancer. However, the mechanisms underlying the anti-tumor properties of CD4^+^ T cells are complex and not yet fully understood. To investigate these mechanisms, we utilized a murine model of tyrosinase-related protein 1 (Trp1)-specific CD4^+^ T cell adoptive transfer therapy for treating melanoma. By employing single-cell RNA sequencing (scRNA-seq), we analyzed the immune cells present in the tumors of mice that received adoptive transfer of Trp1-specific CD4^+^ T cells. Unexpectedly, within the tumor-infiltrating immune cells, the Trp1 CD4^+^ T cell population was relatively small, displaying characteristics indicative of exhaustion. In contrast, the most prominent cell cluster comprised macrophages, expressing high levels of the T cell inhibitory receptor ligand PD-L1 and the pro-inflammatory cytokine IL-1β, suggesting a distinct M1 phenotype. Systemic depletion of macrophages following Trp1 CD4^+^ T cell transfer therapy compromised the antitumor effectiveness and resulted in tumor recurrence. These findings highlight the crucial role of innate macrophages as an effector cell population in Trp1-specific CD4^+^ T cell adoptive cell transfer therapy.

## INTRODUCTION

T lymphocytes are crucial in cancer immunotherapy^1^. Immune checkpoint blockade therapies have shown promising clinical responses in a subset of cancer patients by using monoclonal antibodies to block co-inhibitory molecules such as CTLA-4, PD-1, PD-L1/2, and LAG3^2,3^. Preexisting intratumoral CD8^+^ T cells in metastatic melanoma patients predict response to the PD-1 blockade therapy^4^. In adoptive cell therapy (ACT), autologous T cells are extracted from the patient’s tumors or peripheral blood, modified and amplified in vitro, and then reintroduced into the patient^5^. Adoptively transferred T cells are usually composed of CD8^+^ and CD4^+^ T cells, and the anti-melanoma activity of which is mainly attributed to the cytotoxicity of CD8^+^ T cells^6^. In breast cancer patients, tumor-infiltrating CD8^+^ T cells can predict clinical outcomes^7^. However, the anti-cancer activity of these CD8^+^ T cells can diminish in exhaustion-inducing immunosuppressive tumor microenvironments^8^.

Recently, as the field of immunotherapy has progressed, more studies have discovered an important role of CD4^+^ T cells in antitumor immunity^9^. Whereas CD8^+^ T cells are thought to be primarily responsible for the direct killing of tumor cells, CD4^+^ T cells help by licensing dendritic cells and secreting inflammatory cytokines such as interferon (IFN)-γ and tumor necrosis factor (TNF)-α, which promote CD8^+^ T cell activity. For instance, upon antigen rechallenge, CD4^+^ T cells play an important role in maintaining a memory CD8^+^ T cell population^10^. However, emerging evidence reveals that CD4^+^ T cells can also express key cytotoxic effector molecules such as granzyme B and perforin. Furthermore, the production of IFNγ by these cells is essential for the rejection of murine melanoma ^11^. A recent single-cell RNA sequencing (scRNA-seq) analysis revealed the enrichment of cytotoxic CD4^+^ T cells instead of CD8^+^ T cells in human bladder cancer, and the cytotoxic CD4^+^ T cell gene signature predicts response to anti-PD-L1 therapy^12^. This study establishes the critical role of cytotoxic CD4^+^ T cells in combatting human bladder cancer, in addition to murine melanoma.

To further dissect the complex mechanisms responsible for the activation and tumor-killing ability of the CD4^+^ T cells, we use the well-established murine transgenic gp75/tyrosinase-related protein 1 (Trp1)-specific CD4^+^ T cells as a model. The self-antigen Trp1 expressed in immunologically “cold” melanoma can be recognized by Trp1-specific TCR on MHC class II-restricted T cells. Adoptive transfer of Trp1 CD4^+^ T cells into B16 melanoma-bearing mice has demonstrated remarkable tumor eradication and the development of autoimmunity within a relatively short timeframe of 35-45 days, even in the absence of exogenous cytokine administration or vaccination. The endogenous CD8^+^ T cells, B cells, NK cells, or NKT cells were found to be dispensable for the potent CD4^+^ T cell-mediated antitumor immunity^13^. In this study, we performed a scRNA-seq analysis on tumor-infiltrating immune cells and revealed a critical role of macrophages in the Trp1 CD4^+^ cells-mediated anticancer activity. Our data ought to provide a stronger understanding of the involvement of the innate immune system in CD4^+^ T cell-mediated cancer immunotherapies.

## MATERIALS AND METHODS

### Mice

*B6. Cg-Rag1tm1Mom Tyrp1B-w Tg(Tcra,Tcrb)9Rest/J* TRP1-specific CD4^+^ TCR transgenic (Trp1) mice and *Rag1*^−/–^ mice, both bred on a B6 background, were purchased from Jackson Laboratory and housed at SPF (specific pathogen-free) conditions at Houston Methodist Hospital. The mice were provided with water and a standard laboratory diet and housed in an environment with a 12-hour light/12-hour dark cycle. Temperatures were maintained between 68–79°F, and humidity levels ranged from 30-70%. For all experiments, mice aged eight to twelve weeks were utilized. Randomization of age- and sex-matched mice was performed before their inclusion in experiments. All animal procedures conducted in this study received approval from the Houston Methodist Animal Care Committee, aligning with institutional animal care and use guidelines.

### Tumor cell line

The B16F10 murine melanoma cells were obtained from the American Type Culture Collection (ATCC) and were tested free from mycoplasma and other pathogens. The cells were cultured in Dulbecco’s Modified Eagle Medium (DMEM) supplemented with 10% fetal bovine serum (FBS) and were maintained in a 37°C incubator with 5% CO_2_ supplementation.

### Tumor Inoculation

To establish a solid tumor model in mice, B16F10 melanoma cells were cultured *in vitro*, collected, and suspended in PBS at a concentration of 2.5 x 10^6^ cells/ml. Subsequently, 5 x 10^5^ cells were subcutaneously injected into the right hind limb of *Rag1*^−/–^ mice. This method was chosen because it mimics the natural growth pattern of melanoma tumors and allows for monitoring of tumor growth and progression over time.

### Adoptive transfer of Trp1-specific CD4^+^ T cells to treat melanoma in mice

Ten days after inoculating B16F10 melanoma cells, spleens and lymph nodes were obtained from Trp1 mice. A single-cell suspension was obtained by grinding and filtering the tissues using a 70 μm cell strainer. Red blood cells were lysed using an Ammonium-Chloride-Potassium (ACK) lysis buffer (Thermo Fisher Scientific). Fifty thousand Trp1 CD4^+^ T cells in PBS were adoptively transferred *via* the tail vein into B16F10 melanoma-bearing mice. The no cell transfer control group was injected with an equal volume of PBS.

### Macrophage depletion by Clodronate liposome injection

On the day of Trp1 CD4^+^ T cell transfer, tumor-bearing mice received an intraperitoneal injection of 200 μl of clodronate liposome (Sapphire North America), followed by another 100 μl of clodronate liposome injection four days later. Tumor growth was monitored, and tumor volumes were measured over time.

### Isolating immune cells from tumors and draining lymph nodes

To analyze tumor-infiltrating immune cells, we injected 5 μg of PE-Cy7 dye-conjugated anti-CD45 (Clone 30-F11, BioLegend) monoclonal antibodies (mAb) into tumor-bearing mice *via* the tail vein. Five minutes later, the mice were euthanized, and their tumors were obtained and placed in a PBS solution. The tumors were minced into small pieces and transferred to a 70 μm cell strainer. A syringe plunger was used to mesh the tissue through the strainer, and the resulting cell suspension was transferred to 50 ml centrifuge tubes containing 10 mL of Ficoll-Paque media (Cytiva) to isolate immune cells, according to the manufacturer’s instruction.

To collect immune cells from the draining lymph nodes, the lymph nodes were mechanically disrupted using the back of a 1-ml syringe, filtered through a 70-μm cell strainer, and incubated with ACK lysis buffer to eliminate red blood cells. The cells were then washed twice with cold PBS supplemented with 2% FBS.

### Fluorescence-activated cell sorting

Single-cell suspensions from the tumors and draining lymph nodes were first stained with 1 μL Zombie Aqua (BioLegend) to exclude dead cells. To remove Zombie Aqua, 5 mL of PBS supplemented with 2% FBS was added, and the solution was centrifuged at 600g for 5 minutes. The supernatant was discarded, and the cells were resuspended in 200 μL of PBS supplemented with 2% FBS. Tumor-infiltrating cells were stained with FITC-conjugated anti-mouse CD45 (Clone 30-F11, BioLegend) and PE CD4 (GK1.5, BioLegend) at a dilution of 1:200. Draining lymph node cells were stained with Pacific blue CD45 (Clone 30-F11, BioLegend) at a dilution of 1:200. After staining, the cells were washed with 20 mL of PBS supplemented with 2% FBS, centrifuged at 600 g for 5 minutes, and the supernatant was discarded. Finally, the cells were resuspended in 500 μL of PBS supplemented with 2% FBS. PE-Cy7^−^FITC^+^ CD45^+^ cells were sorted from tumor-infiltrating cells and Pacific Blue^+^ CD45^+^ cells were sorted from draining lymph node cells using an FACSAria cell sorter (BD Biosciences). The sorted cells were then sent to Baylor College of Medicine for single-cell RNA sequencing. The flow cytometry data were processed using the FlowJo v10 software (Tree Star, Inc).

### Single-cell RNA sequencing analysis

CD45^+^ immune cells were isolated from draining lymph nodes and tumors of B16F10-bearing mice treated with Trp1 cells, as well as from tumors from control mice treated with PBS. The Single Cell Gene Expression Library was created using the Chromium Next GEM Single Cell 3’ Reagent Kits v3 by 10x Genomics. In brief, a suspension of single cells, reverse transcription reagents, Gel Beads containing barcoded oligonucleotides, and oil were loaded onto a Chromium controller (10x Genomics) to create single-cell GEMs (Gel Beads-In-Emulsions). This process facilitated the synthesis of full-length cDNA and barcoding for each individual cell. The GEMs were then broken, and the cDNA from each single cell was pooled. Cleanup was performed using Dynabeads MyOne Silane Beads, followed by PCR amplification of the cDNA. The amplified products were fragmented to an optimal size, and end-repair, A-tailing, and adaptor ligation were carried out. The final library was generated through amplification. Library quantification was conducted using the KAPA Library Quantification kit from Roche, and sequencing was executed on a Novaseq 6000 (Illumina).

The raw Illumina sequencing reads were aligned to the reference mouse genome mm10 (Ensembl 93) using Cell Ranger v7.0.0 with default parameters, resulting in the generation of Fastq datasets. Gene quantification was performed as UMI counts using Cell Ranger count, and initial visualization was conducted. The downstream analysis utilized Seurat V4.0.0 on filtered feature counts generated by Cell Ranger count. Low-quality single cells, containing more than 2.5% mitochondrial transcripts or over 50% ribosomal transcripts, were excluded from the analysis. Additionally, genes expressed in fewer than three single cells were omitted. Potential single-cell doublets were identified using DoubletFinder V2.0.3, with an expected doublet rate of 7.5% assuming Poisson statistics. After excluding low-quality and doublet cells, datasets were merged using Seurat. Single-cell gene expression was normalized to the library size and log2-transformed in Seurat. Batch correction was implemented using Harmony V1.0. Feature plots and violin plots were generated using Seurat to visualize the expression patterns of key marker genes across the entire dataset.

### Statistical analysis

Statistical analyses were performed using Two-way ANOVA and log-rank test (Prism 8.0.0, GraphPad Software). Tumor volume was assessed with the equation volume = 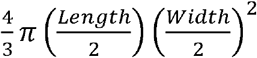. Tests and sample sizes were indicated in the figure legends. A *P* value of <0.05 was considered statistically significant.

## RESULTS

### Adoptively transferred Trp1-specific CD4+ T cells effectively eradicate the established murine melanoma

We employed *Rag1*^−/–^ mice to assess the antitumor activity of Trp1-specific CD4^+^ T (Trp1) cells in the absence of endogenous T cells, B cells, and NKT cells. As is shown in **Figure 1A**, we subcutaneously (S.C.) injected 5 x 10^5^ B16-F10 melanoma cells into the right hind limb of mice to induce melanoma and subsequently monitored tumor growth. When the tumors reached approximately 0.8 cm in diameter (typically 10 days after tumor implantation), we administered a tail vein injection of freshly isolated 50,000 naïve Trp1 CD4^+^ T cells into tumor-bearing mice. The control group received PBS treatment at the same time (**Figure 1A**). Quantification of tumor volume revealed that the tumors in the Trp1 CD4^+^ T cell treatment group had a tumor volume of 0.3876 ± 0.3299 cm^3^, in contrast to a tumor volume of 1.9341 ± 0.5361 cm^3^ in the control group on day 17 post-tumor implantation (**Figure 1B**). The adoptive transfer of 50,000 naïve Trp1 T cells into *Rag1*^D/D^ mice on day 10 post-B16F10 melanoma implantation significantly eradicated the established melanomas, leading to prolonged survival in B16F10 melanoma-bearing mice (**Figure 1B,C**).

**Figure 1.**
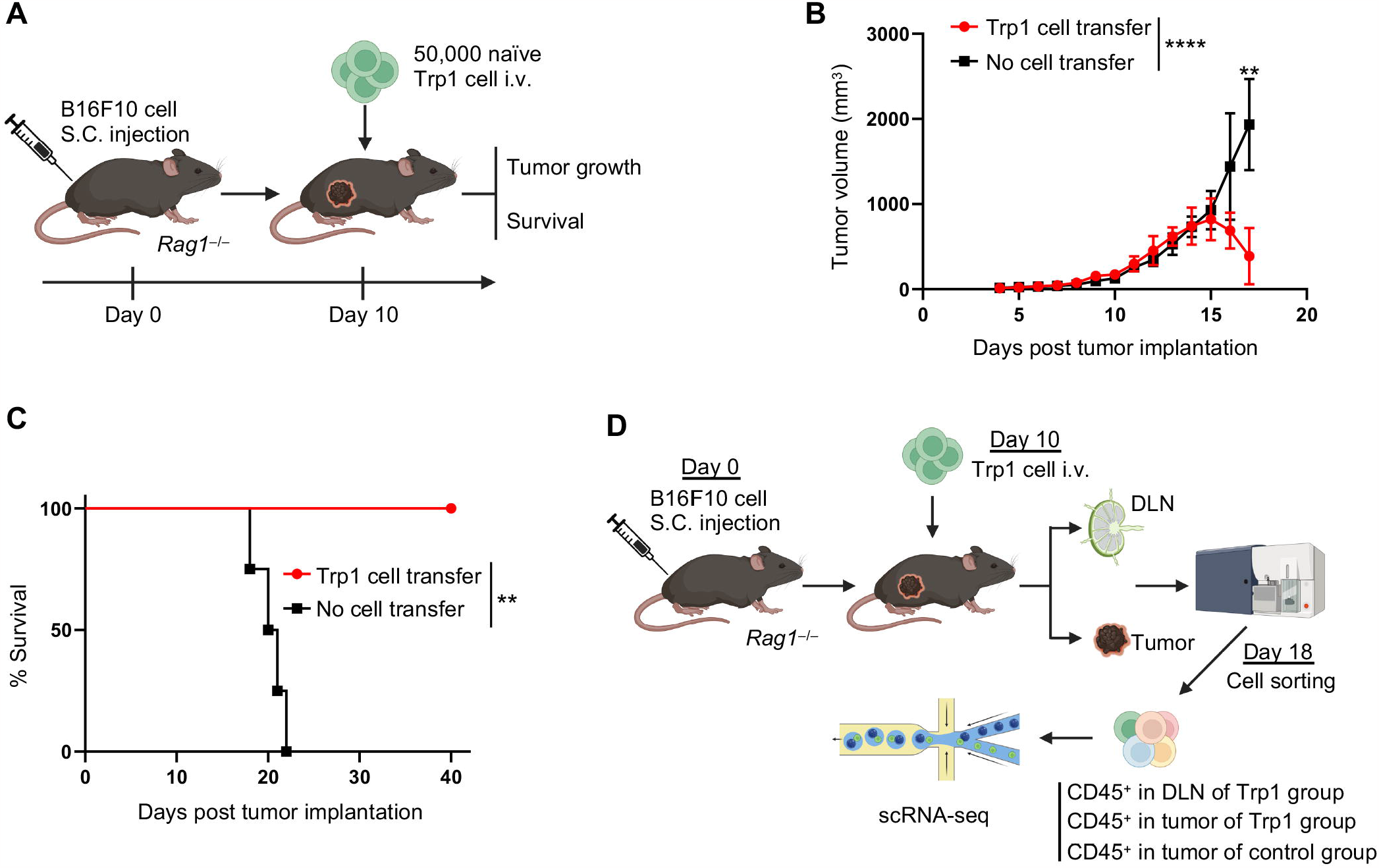
Antitumor activity of transgenic Trp1-specific CD4^+^ T cells in mice. (**A-C**). 5 x 10^5^ B16F10 melanoma cells were subcutaneously injected in *Rag1*^−/–^ mice on day 0. Ten days later, 50,000 naïve Trp1 CD4^+^ T cells were adoptively transferred into these tumor-bearing mice. (**A**) Schematic of the experimental design. (**B**) Tumor volume of B16F10-bearing mice receiving Trp1 cell transfer or PBS (No cell transfer). (**C**) % survival of B16F10-bearing mice receiving Trp1 cell transfer or no cell transfer. (**D**) Schematic of the experimental design for scRNA-seq. In **A** and **D**, schematics of the experimental design were created with BioRender.com. In **B**, data are presented as mean ± SD (n = 5 mice in the Trp1 cell transfer group; n = 4 mice in the no cell transfer group). The *P* values are from the Two-way ANOVA (comparing tumor growth) and unpaired Student’s *t*-test (comparing tumor volumes at day 17 post-tumor implantation). In **C**, the *P* value is from the log-rank test. **, *P* < 0.01; ****, *P* < 0.0001.

To explore the mechanism underlying Trp1 CD4^+^ T cell-mediated tumor regression, we conducted scRNA-seq analysis of CD45^+^ immune cells isolated from the draining lymph nodes and tumors of B16F10-bearing mice treated with Trp1 CD4^+^ T cells, as well as tumors from control mice treated with PBS (**Figure 1D**). **Figure S1** shows the gating strategy for sorting tumor-infiltrating CD45^+^ cells for scRNA-seq analysis. Two different anti-CD45 antibodies were employed to distinguish tumor-infiltrating CD45^+^ cells from circulating CD45^+^ cells. Flow cytometry analysis revealed that Trp1 CD4^+^ T cells can be detected in Trp1 cell transfer group, but not in the no cell transfer control group (**Figure S1**).

### ScRNA-seq analysis of lymph node-resident and tumor-infiltrating immune cells

To further characterize the functional states of individual immune cells and their gene expression profiles, we performed single-cell RNA-sequencing on CD45^+^ immune cells isolated from the draining lymph nodes and the tumors in the Trp1 CD4^+^ T cell treatment group (**Figure 1D**). Clustering analysis identified 21 clusters of immune cells (**Figure 2A, B**). Violin plots were utilized to illustrate the expression profiles of signature genes at the single-cell level. Clusters 1, 3, 5, 7, 8, 10, 11, 12, 13, 17, and 18 exhibited elevated expressions of T cell marker genes, including *Cd3e* and *Trac*, identifying them as T cell clusters (**Figure 2C, S2A**). Cluster 9, 14, and 16 were characterized as NK cells based on the expression of NK cell marker genes such as *Ncr1, Klre1*, and *Klrb1c* (**Figure 2C, S2B**). Among the clusters expressing high levels of the myeloid cell marker *lgtam* (clusters 0, 2, 4, 6, 9, 12, 15, 18, and 20), clusters 0, 4, 15, and 19 prominently expressed *Cd68* and were identified as macrophages (**Figure S3A**). Neutrophils (clusters 2, 6, and 18), basophils (cluster 20), and DCs (cluster 17) were also identified, with a higher abundance in the lymph nodes compared to tumors (**Figure S2C**,**D, S3B**). Notably, the most prevalent population of immune cells in Trp1 CD4^+^ T cell-treated tumors was cluster 0, consisting of PD-1^hi^IL-1^hi^ macrophages, which were scarcely detected in the draining lymph nodes. Conversely, the transferred Trp1 CD4^+^ T cells predominantly resided in the draining lymph nodes, with only a small fraction infiltrating the established tumors and facilitating tumor regression (**Figure 2B,C**, **S2A**). Hence, scRNA-seq analysis revealed distinct immune subsets in the draining lymph nodes and tumor microenvironment following Trp1 cell treatment.

**Figure 2.**
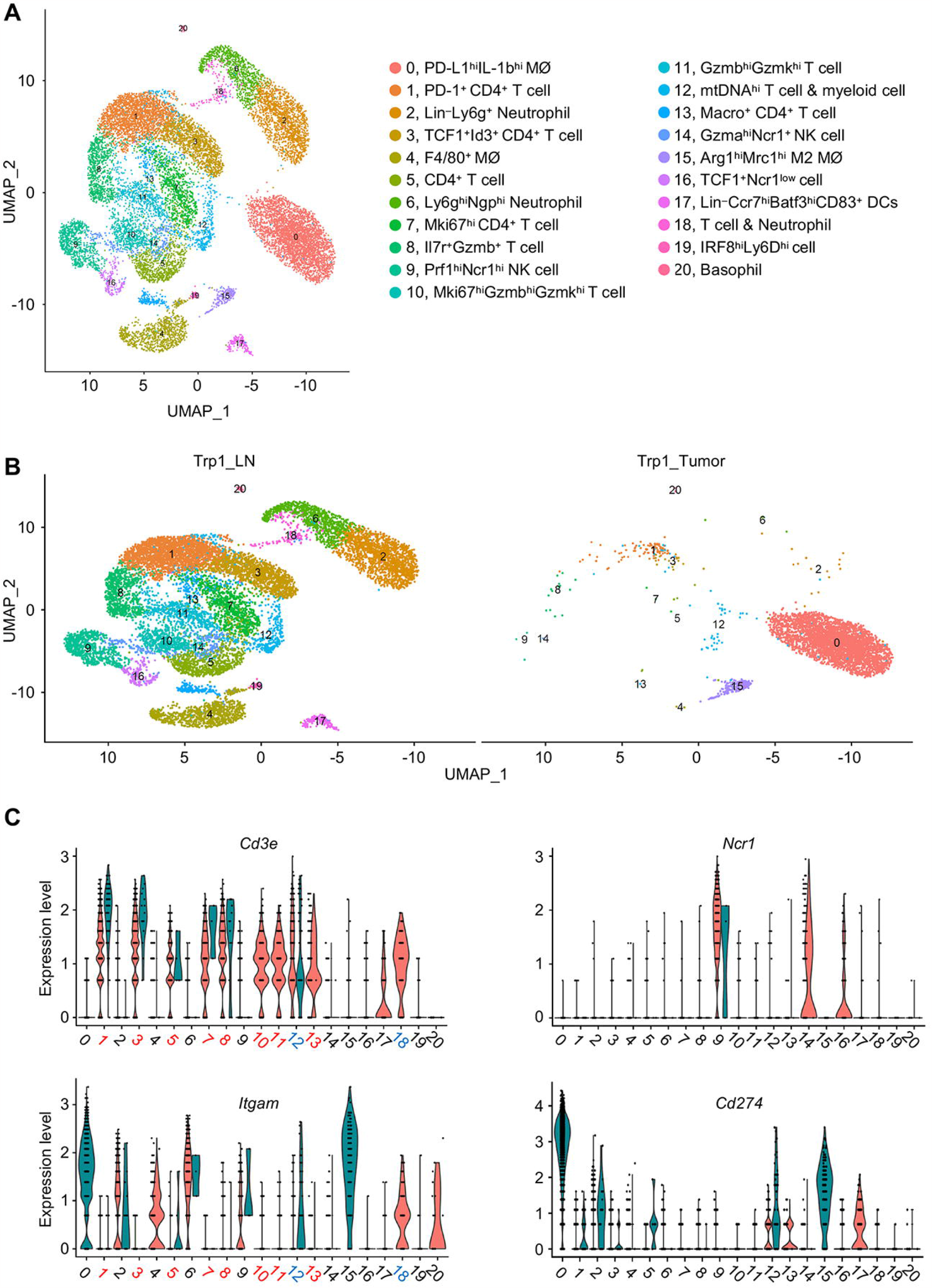
ScRNA-seq reveals different immune cell subsets in the tumors and lymph nodes from B16F10 tumor-bearing mice treated with Trp1 CD4^+^ T cells. (**A-C**). On day 0, *Rag1*^−/–^ mice were subcutaneously injected with 5 x 10^5^ B16F10 melanoma cells. Ten days later, 50,000 naive Trp1 CD4^+^ T cells were adoptively transferred into these mice bearing tumors. On day 18 post-tumor implantation, CD45^+^ cells from tumors (Trp1_Tumor) and draining lymph nodes (Trp1_LN) were sorted for scRNA-seq analysis. (**A)** UMAP projections of the combined scRNA-seq datasets from Trp1_Tumor and Trp1_LN. (**B**) UMAP projections of the split scRNA-seq dataset from Trp1_Tumor or Trp1_LN. (**C**) Violin plots showing the expression of *Cd3e* (T cells), *Ncr1* (NK cells), *Itgam*/CD11b, and *Cd274*/PD-L1 markers.

### Trp1 CD4+ T cells in melanoma exhibit an exhaustion-like phenotype

Clusters 1, 3, 8, and 12 are the main T cell clusters in tumors. These clusters exhibited comparable expression levels of stem-like genes, including *Tcf7, Sell, II7r*, and *Id3*, in comparison to other T cell clusters present in the draining lymph nodes (**Figure 3A**). Furthermore, they displayed higher expression levels of exhaustion markers such as *Pdcd1, Lag3, Havcr2*, and *Nr4a1*, as well as lower expression levels of effector genes including *Prf1, Gzmb, Mki67*, and *Gzmk*, suggesting an exhaustion-like phenotype (**Figure 3B, 3C**). Since Trp1 CD4^+^ T cells primarily resided in the draining lymph nodes and exhibited an exhaustion-like phenotype upon infiltration into the tumor, we speculated that there might be additional T cell-extrinsic mechanisms contributing to tumor regression.

**Figure 3.**
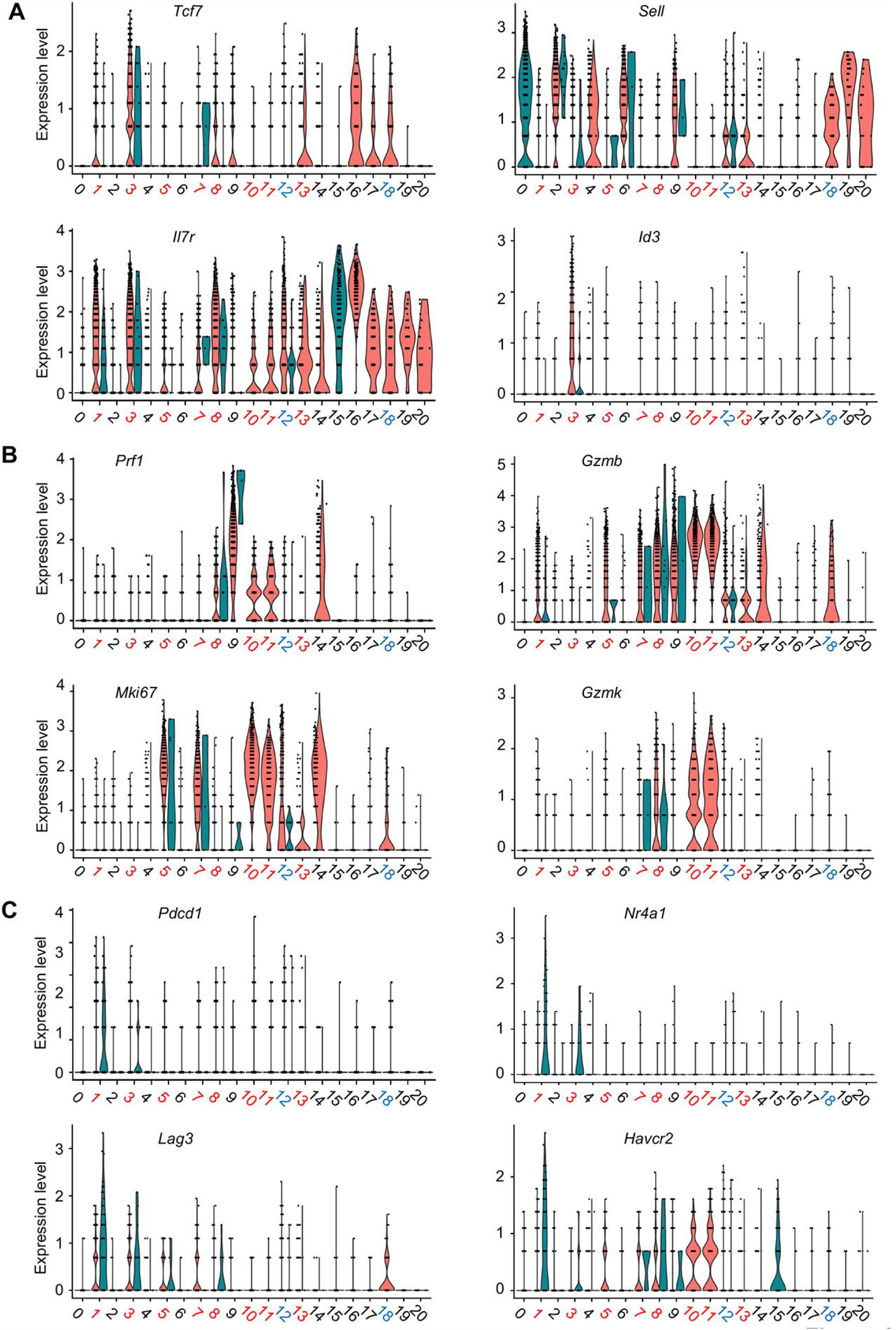
ScRNA-seq reveals distinct cell states of Trp1 cells from tumors and draining lymph nodes in Trp1 cell-treated B16F10 tumor-bearing mice. (**A**) Expression of stemness genes *Tcf7, Sell, Il7r*, and *Id3*. (**B**) Expression of effector genes *Prf1, Gzmb, Mki67*, and *Gzmk*. (**C**) Expression of exhaustion marker genes *Pdcd1, Nr4a1, Lag3*, and *Havcr2* in 21 clusters of immune cells. Cell clusters are indicated on the X-axis, with red-colored numbers representing T cell clusters. The green color halves indicate cells derived from tumors, while the red color halves indicate cells derived from draining lymph nodes.

### PD-L1hiIL-1βhiCD68+ macrophage subset mediates antitumor activity in Trp-1 CD4+ T cell transfer therapy

The scRNA-seq analysis revealed that, in mice treated with Trp1 CD4^+^ T cells, a large population of macrophages were present in the tumors, but not in the lymph nodes. To investigate whether these tumor-infiltrating macrophages were induced by transferred Trp1 CD4^+^ T cells, we compared scRNA-seq results of the tumor-infiltrating immune cells from mice treated with PBS and mice treated with Trp1 CD4^+^ T cells. Shown in **Figure 4A**, scRNA-seq analysis identified 16 clusters of immune cells. As expected, a small population of CD4^+^ T cells, cluster 13, only existed in Trp1 cell-treated tumors, but not in the tumors from the control mice (**Figure 4B**). The control tumors contained a large number of NK cells (cluster 4), H2-A^hi^CCR7^+^ DC (cluster 5), neutrophils (cluster 6), H2-A^hi^ DC (cluster 9), CCR7^hi^ DC (cluster 14), and cDC1 & cDC2 (cluster 15). The large number of macrophages existed in both groups of tumors, however, with different molecular characteristics. For instance, PD-L1^hi^ macrophages (cluster 0 and 2), CCR5^+^CD38^+^ macrophages (cluster 10), and Arg1^hi^Mrc1^hi^ M2 macrophages (cluster 12) were only presented in Trp1 cell-treated tumors. In contrast, control tumors contained F4/80^+^CD38^+^ macrophages (cluster 1), LysM^hi^F4/80^+^ macrophages (cluster 3), LysM^+^F4/80^+^ macrophages (clusters 7 and 8, and CCR5^+^CD38^+^ macrophages (clusters 10 and 11). Violin plots revealed that cluster 13 is comprised of T cells, cluster 4 of NK cells, and cluster 6 of neutrophils. Cluster 0 and 2 macrophages, exclusively identified in tumors treated with Trp1 cells, expressed high levels of *Cd274*/PD-L1 (**Figure 4C**). Therefore, the major cell cluster observed in the regressed tumor was identified as PD-L1^hi^IL-1b^hi^CD68^+^ macrophages, implying their potential role in mediating antitumor activity in Trp1 CD4^+^ T cell therapy.

**Figure 4.**
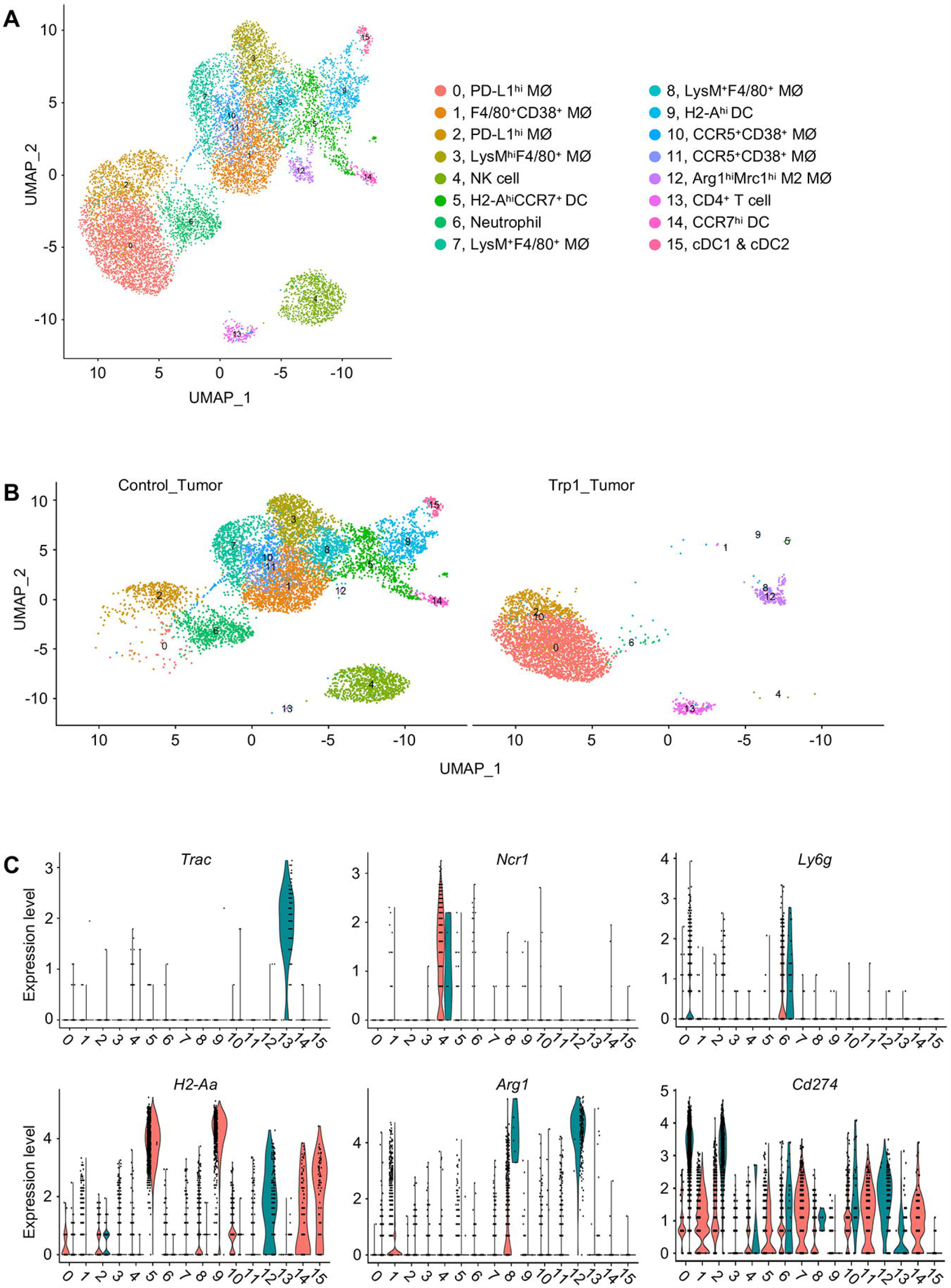
Comparison of gene expression pattern of tumor-infiltrating immune cells between Trp1 adoptive transfer therapy group and control group. **(A-C)** On day 0, *Rag1*^−/–^ mice were subcutaneously injected with 5 x 10^5^ B16F10 melanoma cells. Ten days later, 50,000 naïve Trp1 CD4^+^ T cells (Trp1_Tumor) or PBS control (Control_tumor) were adoptively transferred into these tumor-bearing mice. On day 18 post-tumor implantation, CD45^+^ cells from tumors were sorted for scRNA-seq analysis. (**A)** UMAP projections of the combined scRNA-seq datasets from Trp1_Tumor and Control_tumor. (**B**) UMAP projections of the split scRNA-seq dataset from Trp1_Tumor or Control_tumor. (**C**) Violin plots showing the expression of *Trac, Ncr1, Ly6g, H2-Aa, Arg1*, and *Cd274*.

### Systemic macrophage depletion following adoptive transfer of Trp1 CD4+ T cells in melanoma-bearing mice led to tumor recurrence

It has been demonstrated previously that the expression of *Cd274*/PD-L1 in macrophages indicates a tumor-associated macrophage (TAM) phenotype, which is responsible for promoting an immunosuppressive tumor microenvironment^14^. However, the presence of *Il1b* also suggests a role in cytokine production, alluding to a classically activated M1 phenotype^12^. To determine the role of the PD-L1^hi^IL-1b^hi^CD38^+^ macrophages, we depleted macrophages in mice with clodronate liposomes, following adoptive transfer of Trp1 CD4^+^ T cells into B16 melanoma-bearing mice (**Figure 5A**). Depletion of macrophage resulted in a significant reduction in the antitumor activity of adoptive transfer therapy, as evidenced by tumor recurrence by day 50. In contrast, in the non-depletion group, mice exhibited complete tumor remission by day 27 (**Figure 5B**). Hence, systemic macrophage depletion following the adoptive transfer of Trp1 CD4^+^ T cells in melanoma-bearing mice led to tumor recurrence.

**Figure 5.**
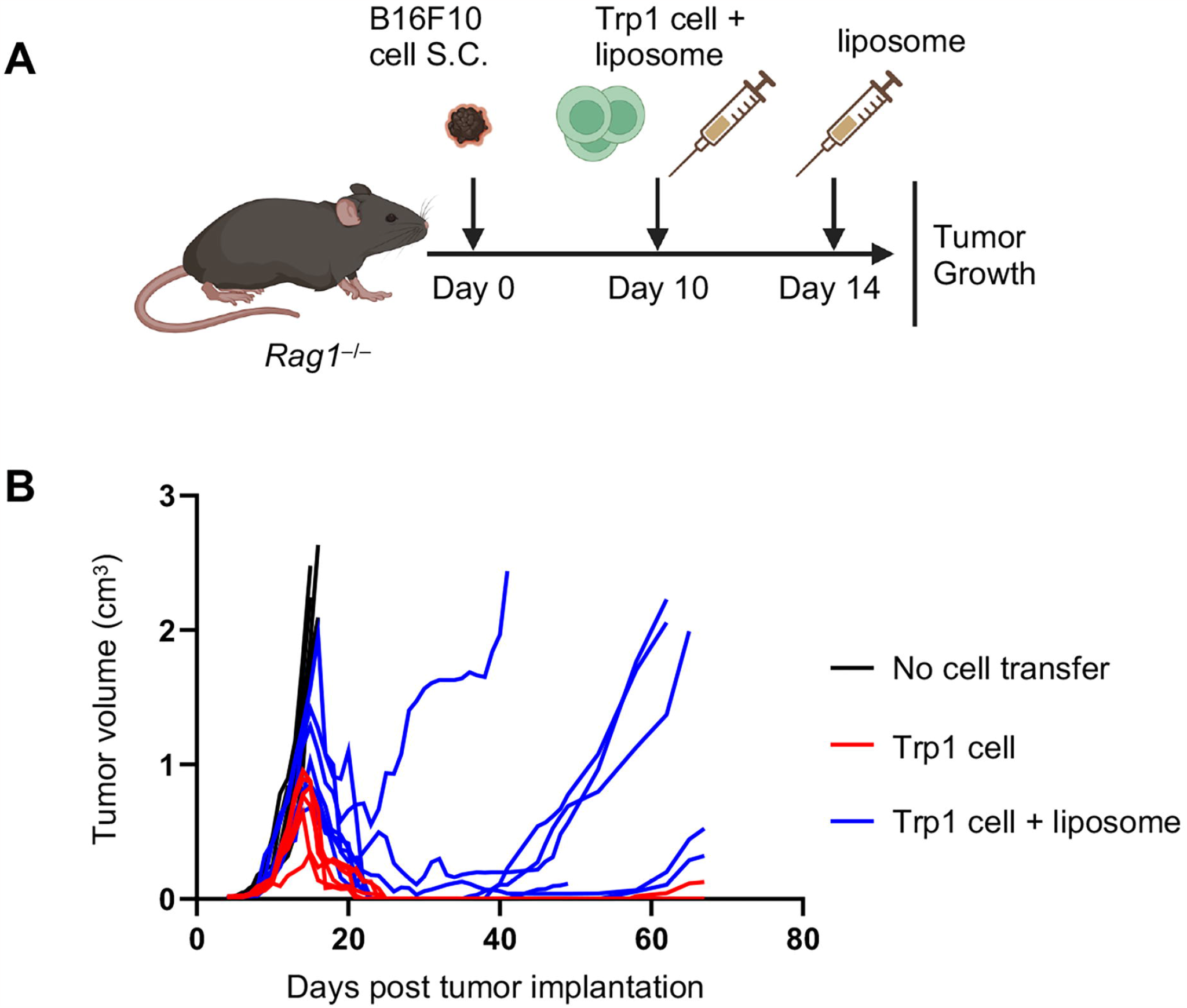
Depletion of macrophages with clodronate liposomes leads to tumor recurrence. (**A,B**) On day 0, *Rag1*^−/–^ mice were subcutaneously injected with 5 x 10^5^ B16F10 melanoma cells. Ten days later, these mice were treated with 50,000 naïve Trp1 CD4^+^ T cells alone, 50,000 naïve Trp1 CD4^+^ T cells plus 200 μl liposome, or PBS (no cell transfer). Four days later, in the group treated with 50,000 Trp1 CD4^+^ T cells plus 200 μl liposome, mice were treated with 100 μl liposome. (**A**) Schematic of the experimental design, created with BioRender.com. (**B**) Tumor volumes of mice treated with Trp1 CD4^+^ T cells alone, Trp1 CD4^+^ T cells plus 200 μl liposome, or PBS (no cell transfer). Data are presented as mean ± SD (n =5-7 mice per group).

## DISCUSSION

Emerging evidence highlights a critical role of CD4^+^ T cells in cancer immunotherapy. Using the transgenic CD4^+^ T cells that recognize the melanoma self-antigen Trp1, we confirmed that the Trp1-specific CD4^+^ T cells were highly effective in eradicating large subcutaneous B16 melanoma in mice. However, in a previous report, the anti-melanoma efficacy of the CD4^+^ T cells was largely attributed to a lymphopenic microenvironment, which induced the differentiation of Trp1 CD4^+^ T cells into a cytotoxic Th1 subset^15^. It was also shown that IFNγ secreted by Trp1 CD4^+^ T cells was critical for tumor clearance, whereas TNFα, endogenous T, B, NK, or NKT cells did not play a significant role^11,15^.

In this study, our findings suggest that the antitumor activity of Trp1 CD4^+^ T cells cannot be solely attributed to their cytotoxic properties. Following adoptive transfer therapy, Trp1 CD4^+^ T cells constituted only a small fraction of the tumor-infiltrating immune cell population. Instead, our analysis utilizing scRNA-seq unveiled a substantial presence of TAMs. TAMs are conventionally classified into two distinct subsets, each exerting opposing roles in cancer development: classically activated M1 and alternatively activated M2 macrophages^16^. M1 macrophages, stimulated by IFNγ and LPS, demonstrate tumoricidal activity, employing the production of reactive oxygen species and proinflammatory cytokines including IL-1β, IL-6, and TNFα. Conversely, M2 macrophages, activated by IL4 or IL13, promote tumorigenesis and cancer progression by generating anti-inflammatory cytokines like IL-10, IL-13, and TGF-β.

Based on our scRNA-seq analysis, we identified two distinct sub-populations of macrophages within the melanoma microenvironment following Trp1 CD4^+^ cell treatment: cluster 0 and cluster 15. Notably, macrophages in the much smaller cluster 15 exhibited the expression of Arg1 and Mrc1, which are recognized markers associated with M2 macrophages. Conversely, the macrophages in the predominant cluster 0 displayed elevated levels of PD-L1 and the inflammatory cytokine IL-1β, indicating their resemblance to M1 macrophages. Intriguingly, the systemic depletion of macrophages abolished the anti-cancer activity of adoptive transferred Trp1 CD4^+^ T cells in melanoma-bearing mice, providing compelling evidence for the crucial role of tumor-infiltrating M1 macrophages in mediating anti-cancer effects.

The substantial population of cluster 0 macrophages characterized by high expression of PD-L1 and IL-1b was exclusively observed in the tumors treated with Trp1 CD4^+^ T cells, while absent in the control tumors treated with PBS, indicating that the presence of cluster 0 macrophages was induced specifically by Trp1 CD4^+^ T cells. This finding aligns with earlier studies demonstrating that Trp1 CD4^+^ T cells mediate melanoma eradication in an IFNγ-dependent manner, with IFNγ being a key factor in activating M1 macrophages^15,17^. Therefore, our results support the notion that the presence of Trp1 CD4^+^ T cells leads to the induction of cluster 0 macrophages with an M1-like phenotype through the production of IFNγ.

Our findings are consistent with a recent study indicating that the elimination of melanoma by Trp1 CD4^+^ T cells in mice necessitates the involvement of innate immune cells, specifically neutrophils^18^. However, there are notable variations in the experimental approaches employed. Firstly, in the aforementioned study, the administration of an anti-OX40 antibody was combined with Trp1 CD4^+^ T cells, whereas our current study did not utilize the anti-OX40 antibody. Indeed, the omission of an anti-OX40 antibody resulted in a significantly lower presence of tumor-infiltrating neutrophils^18^. Secondly, while we employed *Rag1*^−/–^ mice, the previous study utilized immune-competent wild-type B6 mice that were subjected to cyclophosphamide (CTX) treatment before the adoptive transfer of Trp1 CD4^+^ T cells. These differences in experimental design may have contributed to the recruitment of distinct types of innate immune cells. Nonetheless, both investigations demonstrated the crucial role played by innate immune cells in the regression of tumors induced by Trp1 CD4^+^ T cells.

Collectively, these results provide support for a two-step model underlying the eradication of solid tumors by tumor-reactive CD4^+^ T cells. Firstly, Trp1 CD4^+^ T cells infiltrate melanoma and initiate cancer cell death. Secondly, these tumor-killing T cells generate inflammatory cytokines, such as IFNγ, which in turn recruit and activate innate immune cells such as macrophages and/or neutrophils. These activated innate immune cells play a crucial role in the eradication of solid tumors. Our results indicate that both cytotoxic Trp1 CD4^+^ T cells and innate macrophages/neutrophils act as effectors in tumor regression, and their concerted efforts contribute to the rejection of tumors in B16 melanoma-bearing mice.

## ACKNOWLEDGMENTS

The authors thank the Houston Methodist Research Institute and the Single Cell Genomics Core at the Baylor College of Medicine for their generous support throughout this project. Additionally, we express sincere appreciation to Dr. Yulin Dai at the University of Texas Health Science Center at Houston for his invaluable guidance on the analysis of scRNA-seq data.

## DATA AVAILABILITY

The scRNA-seq data have been deposited in the Gene Expression Omnibus under accession number GSE250511. All data generated or analyzed during this study are included in this manuscript (and its supplemental information files).

## AUTHOR CONTRIBUTIONS

A.H., D.Z., and W.C. designed the study and wrote the manuscript. A.H. and D.Z. performed core experimental work and data analysis. W.C. supervised the study. All authors reviewed and approved the manuscript.

## REFERENCES

1 Waldman, A. D., Fritz, J. M. & Lenardo, M. J. A guide to cancer immunotherapy: from T cell basic science to clinical practice. Nat Rev Immunol 20, 651–668, doi:10.1038/s41577-020-0306-5 (2020).

2 Morad, G., Helmink, B. A., Sharma, P. & Wargo, J. A. Hallmarks of response, resistance, and toxicity to immune checkpoint blockade. Cell 184, 5309–5337, doi:10.1016/j.cell.2021.09.020 (2021).

3 Tawbi, H. A. et al. Relatlimab and Nivolumab versus Nivolumab in Untreated Advanced Melanoma. N Engl J Med 386, 24–34, doi:10.1056/NEJMoa2109970 (2022).

4 Tumeh, P. C. et al. PD-1 blockade induces responses by inhibiting adaptive immune resistance. Nature 515, 568–571, doi:10.1038/nature13954 (2014).

5 Rosenberg, S. A. & Restifo, N. P. Adoptive cell transfer as personalized immunotherapy for human cancer. Science 348, 62–68, doi:10.1126/science.aaa4967 (2015).

6 Radvanyi, L. G. et al. Specific lymphocyte subsets predict response to adoptive cell therapy using expanded autologous tumor-infiltrating lymphocytes in metastatic melanoma patients. Clin Cancer Res 18, 6758–6770, doi:10.1158/1078-0432.CCR-12-1177 (2012).

7 Mahmoud, S. M. et al. Tumor-infiltrating CD8+ lymphocytes predict clinical outcome in breast cancer. J Clin Oncol 29, 1949–1955, doi:10.1200/JCO.2010.30.5037 (2011).

8 Cheng, H., Ma, K., Zhang, L. & Li, G. The tumor microenvironment shapes the molecular characteristics of exhausted CD8(+) T cells. Cancer Lett 506, 55–66, doi:10.1016/j.canlet.2021.02.013 (2021).

9 Tay, R. E., Richardson, E. K. & Toh, H. C. Revisiting the role of CD4(+) T cells in cancer immunotherapy-new insights into old paradigms. Cancer Gene Ther 28, 5–17, doi:10.1038/s41417-020-0183-x (2021).

10 Janssen, E. M. et al. CD4+ T cells are required for secondary expansion and memory in CD8+ T lymphocytes. Nature 421, 852–856, doi:10.1038/nature01441 (2003).

11 Quezada, S. A. et al. Tumor-reactive CD4(+) T cells develop cytotoxic activity and eradicate large established melanoma after transfer into lymphopenic hosts. J Exp Med 207, 637–650, doi:10.1084/jem.20091918 (2010).

12 Oh, D. Y. et al. Intratumoral CD4(+) T Cells Mediate Anti-tumor Cytotoxicity in Human Bladder Cancer. Cell 181, 1612–1625 e1613, doi:10.1016/j.cell.2020.05.017 (2020).

13 Muranski, P. et al. Tumor-specific Th17-polarized cells eradicate large established melanoma. Blood 112, 362–373, doi:10.1182/blood-2007-11-120998 (2008).

14 Sumitomo, R. et al. PD-L1 expression on tumor-infiltrating immune cells is highly associated with M2 TAM and aggressive malignant potential in patients with resected non-small cell lung cancer. Lung Cancer 136, 136–144, doi:10.1016/j.lungcan.2019.08.023 (2019).

15 Xie, Y. et al. Naive tumor-specific CD4(+) T cells differentiated in vivo eradicate established melanoma. J Exp Med 207, 651–667, doi:10.1084/jem.20091921 (2010).

16 Poh, A. R. & Ernst, M. Targeting Macrophages in Cancer: From Bench to Bedside. Front Oncol 8, 49, doi:10.3389/fonc.2018.00049 (2018).

17 Nathan, C. F., Murray, H. W., Wiebe, M. E. & Rubin, B. Y. Identification of interferongamma as the lymphokine that activates human macrophage oxidative metabolism and antimicrobial activity. J Exp Med 158, 670–689, doi:10.1084/jem.158.3.670 (1983).

18 Hirschhorn, D. et al. T cell immunotherapies engage neutrophils to eliminate tumor antigen escape variants. Cell 186, 1432–1447 e1417, doi:10.1016/j.cell.2023.03.007 (2023).

