## Supplemental Figures for "Macrophages as Key Mediators of Trp-1-Specific CD4^+^ T Cell Therapy for Murine Melanoma"

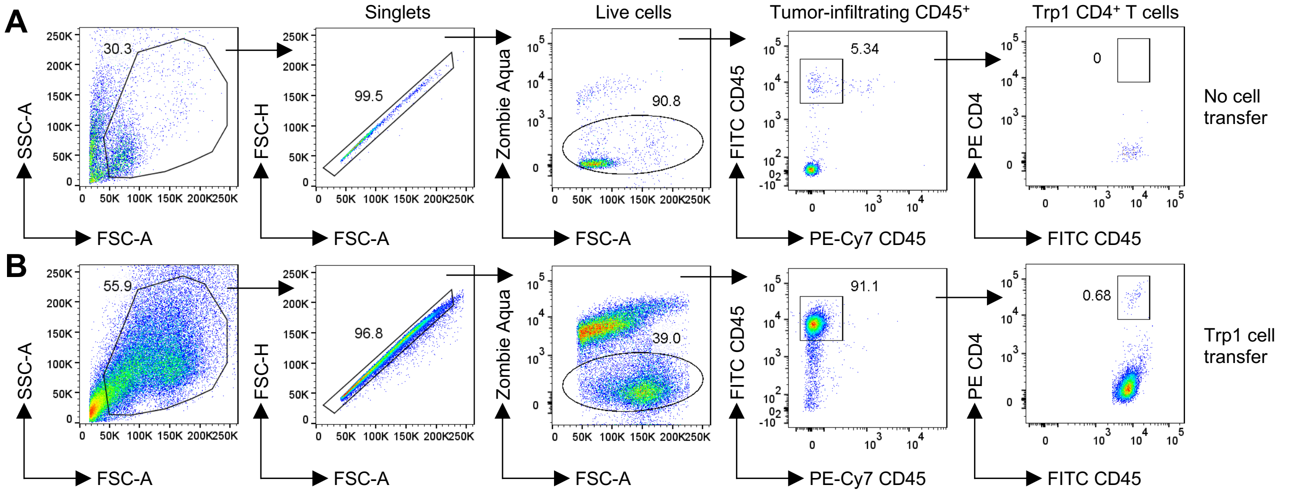


**Figure S1. Gating strategies for cell sorting of tumor-infiltrating immune cells.** At day 0, 5 x $10^5$ B16F10 melanoma cells were subcutaneously injected in *Rag1*$^{-/-}$ mice. Ten days later, PBS vehicle (**A**) or 50,000 naïve Trp1 $CD4^+$ T cells (**B**) were adoptively transferred into these tumor-bearing mice. On day 18 post-tumor implantation, $CD45^+$ cells from tumors were sorted for scRNA-seq analysis.

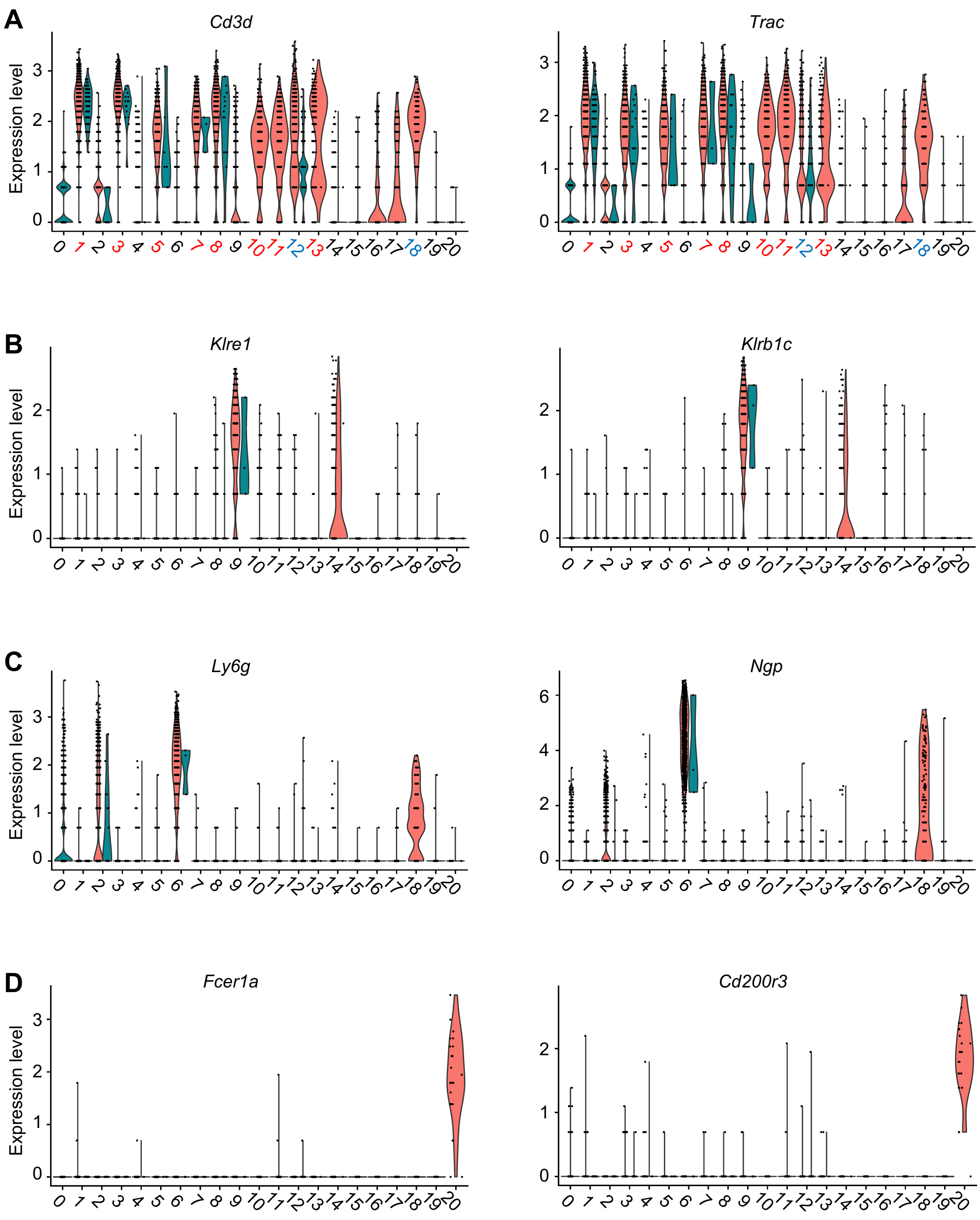


**Figure S2, related to Figure 2**. (**A**) genes expressed in T cells; (**B**) genes expressed in NK cells; (**C**) genes expressed in Neutrophils; (**D**) genes expressed in Basophils.

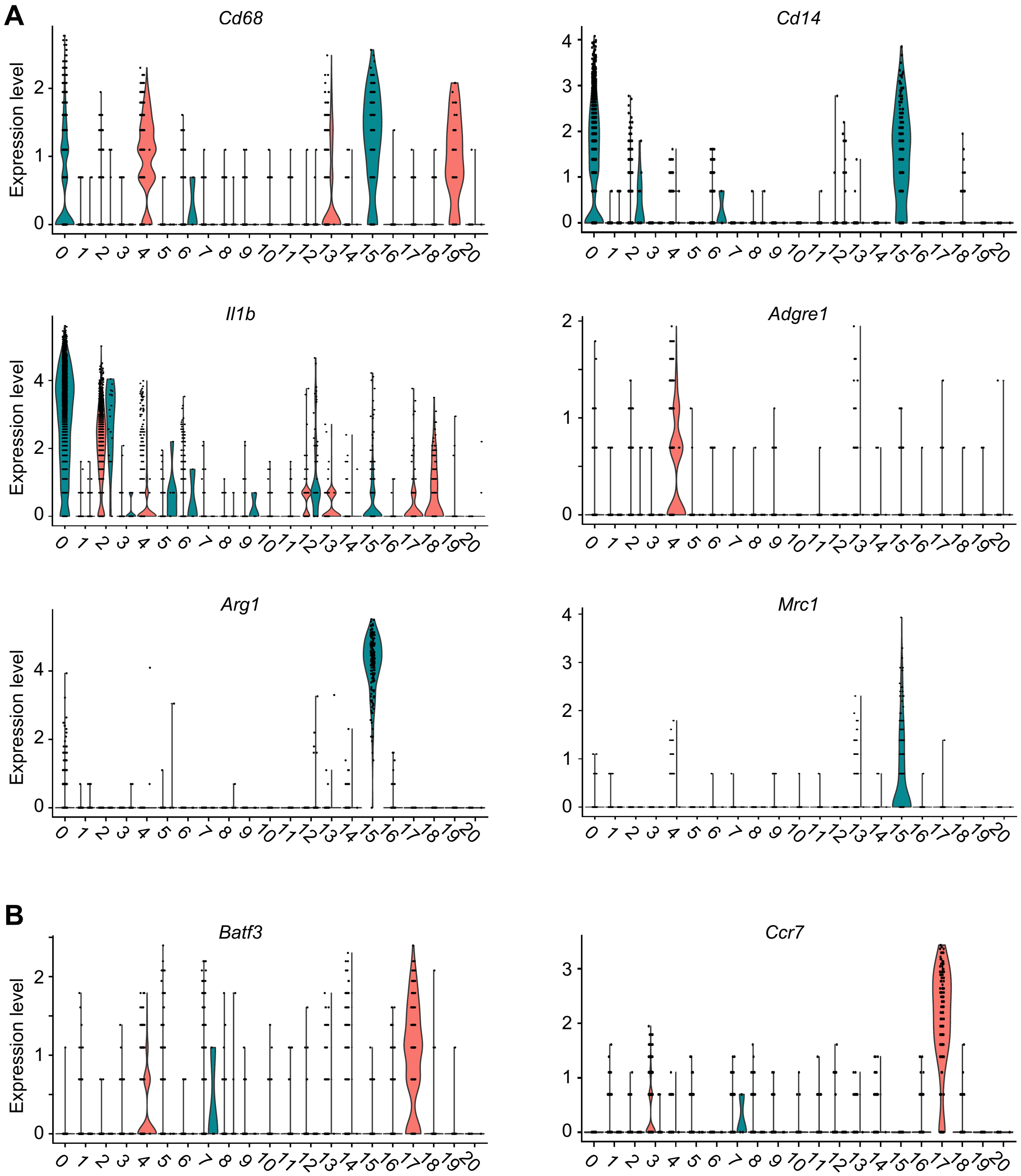


**Figure S3, related to Figure 2**. (**A**) genes expressed in macrophage subsets; (**B**) genes expressed in DCs.
